# Losartan Increases Survival of the *Fbn1^mgR/mgR^* Mouse Model of Marfan Syndrome in an Age-Dependent Manner

**DOI:** 10.1101/2021.02.19.429438

**Authors:** Jeffrey D. Smith, Jeff Z. Chen, Rebecca Phillips, Alan Daugherty, Mary B. Sheppard

**Affiliations:** Saha Cardiovascular Research Center, University of Kentucky, Lexington, KY; Saha Aortic Center, University of Kentucky, Lexington, KY; Department of Surgery, University of Kentucky, Lexington, KY; Department of Physiology, University of Kentucky, Lexington, KY; Department of Family Medicine, University of Kentucky, Lexington, KY

## Abstract

Clinical trials investigating angiotensin receptor blockers (ARB) for attenuation of thoracic aortic aneurysm in people with Marfan syndrome have demonstrated variable efficacy. The primary objective of this study was to determine whether the age of mice at the time of losartan initiation affected mortality in fibrillin-1 hypomorphic (Fbn1mgR/mgR) mice. Male (n=40) and female (n=28) Fbn1mgR/mgR mice were randomized to receive losartan in drinking water (0.6 g/L) starting at either 24 or 50 days of age. Controls included Fbn1mgR/mgR mice (20M, 14F) and wild type (15M, 15F) littermates who were not administered the drug. Mortality of Fbn1mgR/mgR males receiving losartan at postnatal day 24 (P24) was not different from wild type controls (p=0.138). Survival of Fbn1mgR/mgR males administered losartan at P50 was not different compared to Fbn1mgR/mgR males receiving no drug (p=0.194) and decreased compared to wild type mice (p=0.002). Survival analysis after P50 demonstrated increased survival of Fbn1mgR/mgR males administered losartan at P50 compared to Fbn1mgR/mgR mice receiving no drug (p=0.017). Age is a critical variable that affects the therapeutic efficacy of losartan in male Fbn1mgR/mgR mice. Since overall mortality in female Fbn1mgR/mgR mice was lower than in male Fbn1mgR/mgR mice, a survival benefit with losartan was not detected in females.

---

Clinical trials to determine the benefits of angiotensin receptor blockers (ARBs) in treatment of thoracic aortic disease in Marfan syndrome have demonstrated variable efficacy (1). We hypothesized that a critical variable influencing the results was age heterogeneity between trials. To test this hypothesis, we determined the effects of age of losartan initiation on survival in fibrillin-1 hypomorphic (*Fbn1^mgR/mgR^*) mice, a mouse model of Marfan syndrome that develops severe thoracic aortic aneurysms and death due to aortic rupture.

Male (n=40) and female (n=28) *Fbn1^mgR/mgR^* mice were randomized to receive losartan in drinking water (0.6 g/L) starting at either postnatal day 24 (P24) or 50 (P50). Data derived from these mice was compared to littermate *Fbn1^mgR/mgR^* (20 males, 14 females) and wild type (15 males, 15 females) mice that were given vehicle only. All mice were terminated at 20 weeks of age. Mice that died during the study were necropsied to determine cause of death.

Survival of *Fbn1^mgR/mgR^* males receiving losartan at P24 was increased compared to *Fbn1^mgR/mg^*^R^ males receiving no drug (p<0.001, log-rank survival) and not different from wild type male mice (p=0.138). Survival of *Fbn1^mgR/mgR^* males administered losartan at P50 was not different compared to *Fbn1^mgR/mgR^* males receiving no drug (p=0.194) and decreased compared to wild type mice (p=0.002; Figure A). However, several *Fbn1^mgR/mgR^* mice died prior to initiation of losartan at P50. To control for these effects, we performed a survival analysis starting at P50, which demonstrated increased survival of *Fbn1^mgR/mgR^* males administered losartan at P50 compared to *Fbn1^mgR/mgR^* mice receiving no drug (p=0.017; Figure B). Significant differences between groups did not change when survival analysis included deaths due to all-causes.

Our findings demonstrate that age is a critical variable affecting the therapeutic efficacy of losartan in male *Fbn1^mgR/mgR^* mice. Analysis of data reviewed by Milewicz and Ramirez (2) shows that the survival benefit in male *Fbn1^mgR/mgR^* mice decreases as age of losartan initiation increases. Human trials have also demonstrated a greater efficacy of therapy at younger ages. Subanalysis of the Pediatric Heart Network trial demonstrated that a younger age of losartan initiation (male less than 16 years old and female less than 15) was associated with a greater decrease in aortic-root z scores over time with both losartan (p=0.002) and atenolol (p<0.001) (1).

One significant benefit of this study is the ability to detect significant differences in the primary endpoint of survival. The meta-analysis by Al-Abcha et al. (3) found that aortic dilation was attenuated in patients with Marfan syndrome receiving ARB, but no statistically significant difference in the number of clinical events were observed. A median follow-up period of 8 years was necessary for van Andel et al. to demonstrate that losartan administration reduced number the of deaths compared to the control group (0 vs. 5, p=0.14). Based on our studies in mouse, we predict that the survival benefit of ARB therapy in people with Marfan syndrome will become even more evident over time.

Our study also demonstrated that survival was significantly lower in male versus female *Fbn1^mgR/mgR^* mice (p=0.004), highlighting the impact of sex on disease severity in Marfan syndrome. This sexual dimorphism has been described previously in other Marfan syndrome mouse models and human studies (4). Since survival was not different between *Fbn1^mgR/mgR^* and wild type female mice during the length of our study, it was not possible to detect a survival benefit with losartan. Future studies will need to extend the study duration in order to detect a possible survival benefit of losartan administration in female *Fbn1^mgR/mgR^* mice.

Overall, our results suggest that the earlier ARB therapy is initiated in patients with Marfan syndrome, the greater benefit is likely to be realized. This work was supported by the National Institutes of Health grants K01-HL149984 (Sheppard) and UL1-TR001998.

---

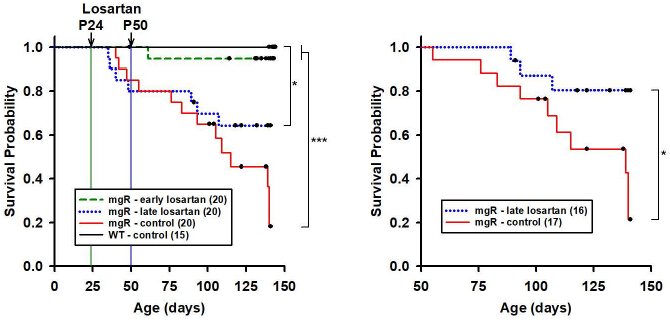
Death by aortic rupture is reduced by losartan administered at P24 versus P50 or controls (left). Losartan at P50 improves survival versus controls (right;* p<0.05, *** p<0.001 by log rank analysis). Censored data include non-rupture deaths and humane endpoints.

## Notes

### Competing Interest Statement

The authors have declared no competing interest.

